# CHD3 regulates BMP signalling response during cranial neural crest cell specification

**DOI:** 10.1101/2025.02.14.638260

**Authors:** Zoe H. Mitchell, Joery den Hoed, Willemijn Claassen, Martina Demurtas, Laura Deelen, Philippe M. Campeau, Karen Liu, Simon E. Fisher, Marco Trizzino

## Abstract

CHD3 is a component of the NuRD chromatin remodeling complex. Pathogenic *CHD3* variants cause Snijders Blok-Campeau Syndrome, a neurodevelopmental disorder with variable features including developmental delays, intellectual disability, speech/language difficulties, and craniofacial anomalies. To unveil the role of CHD3 in craniofacial development, we differentiated *CHD3*-KO induced pluripotent stem cells into cranial neural crest cells (CNCCs). CHD3 expression is low in wild-type iPSCs and neuroectoderm, but upregulated during CNCC specification, where it opens the chromatin at BMP-responsive enhancers, to allow binding of DLX5 and other factors. CHD3 loss leads to repression of BMP target genes and an imbalance between BMP and Wnt signalling, ultimately resulting in aberrant mesodermal fate. Consequently, CNCC specification fails, replaced by early-mesoderm identity, which can be partially rescued by titrating Wnt levels. Our findings highlight a novel role for CHD3 as a pivotal regulator of BMP signalling, essential for proper neural crest specification and craniofacial development.

## INTRODUCTION

Human development is a highly complex process which depends upon the precise regulation of gene expression to determine cell fates. This regulation relies on a series of epigenetic mechanisms, including histone modifications, DNA methylation, and chromatin remodeling.^1,2^ One key chromatin regulator is the nucleosome remodeling and deacetylase (NuRD) complex. The NuRD complex is an ATP-dependent complex which possesses both histone deacetylation and nucleosome remodeling activity.^3–7^ While NuRD was initially thought to act as a repressor of gene expression^3–5^, recent evidence has demonstrated that this complex is able to both repress and activate transcription of target genes through the reorganisation of nucleosome structure.^8–10^ NuRD mediated regulation of gene expression is thought to play a key role during development^11,12^ and is driven by the nucleosome remodeling activity of the complex, which is provided by one of three mutually exclusive CHD subunits: CHD3, CHD4 or CHD5.^10^

The CHD proteins contain an ATPase/helicase domain and a chromodomain motif, which enable the alteration of chromatin structure.^13^ Each NuRD complex only harbours a single CHD protein, with different CHD-NuRD complexes displaying distinct functions and targeting distinct sets of genes.^14^ It has further been suggested that different NuRD configurations may provide time-, tissue- and context-dependent function.^15–17^ For example, a study on mouse cortical development showed that different CHDs were incorporated in the NuRD complex at different developmental stages, with each stage-specific NuRD complex displaying distinct functions.^18^ Given their unique roles, it is perhaps unsurprising that loss or mutation of each one of the CHD proteins results in specific neurodevelopmental disorders.

One example is Snijders Blok-Campeau syndrome, a rare, autosomal dominant, neurodevelopmental disorder resulting exclusively from pathogenic variants within *CHD3*^19^, with affected individuals presenting with a variety of variants, including heterozygous missense variants within the ATPase/helicase domain, and, less frequently, heterozygous loss of function variants.^19–21^ It has been hypothesized that the missense variants may alter the chromatin remodeling ability of CHD3, which could represent a potential pathogenic mechanism behind Snijders Blok-Campeau syndrome.^19^

Affected individuals present with a broad and variable phenotype including different degrees of intellectual disability, impaired speech and language, and macrocephaly^19–21^, along with distinct facial anomalies including a broad, bossed forehead, widely spaced and deep-set eyes, narrow palpebral fissures, midface hypoplasia and low-set ears.^19–21^ So far, only two individuals have been identified with a potential pathogenic *CHD3* variant in both copies of the gene; a homozygous in frame insertion (c.5384_5389dup; p.Arg1796_Phe1797insTrpArg).^22^ These individuals were reported to display a more severe phenotype than cases carrying heterozygous variants, including more distinct facial dysmorphism and severe intellectual disability.^22^ The distinct facial phenotype observed in individuals with pathogenic *CHD3* variants suggests that this NuRD subunit may play an important role in craniofacial development, but this has not been investigated so far.

Craniofacial development is underpinned by the cranial neural crest cells (CNCCs), which constitute an embryonic multipotent cell type from which the bones, cartilage and connective tissues of the face are formed.^23^ CNCCs are generated in the dorsal portion of the neural tube, at the border between the neural plate and the non-neural ectoderm.^24^ Following neural crest induction, these cells undergo epithelial-to-mesenchymal transition (EMT) and subsequently migrate and populate the relevant regions of the developing embryo, where they differentiate into different derivatives, including the craniofacial bones and cartilage.^25^

The process of CNCC specification and formation is complex, and requires the coordinated activity of multiple signalling pathways, key among which are the BMP, and Wnt pathways.^26^ The BMP proteins bind BMP receptors to activate SMAD proteins, which enter the nucleus to trigger specific gene expression programs.^27^ This is mediated by specific BMP-responsive transcription factors, including paralogs of the DLX and MSX families during patterning of the facial mesenchyme.^28–30^ On the other hand, Wnt signalling is required at multiple stages, with roles in neural crest induction, specification, and subsequent migration and differentiation. Wnt ligands bind to Frizzled receptors, allowing β-catenin to enter the nucleus and activate transcription^31^. A finely tuned balance of Wnt, BMP, and FGF signalling is required throughout craniofacial development to enable neural crest induction, specification, migration and subsequent cell fate determination^26,32–39^. We hypothesise that factors such as CHD3 establish appropriate CNCC-chromatin state allowing synchronisation of signalling pathways during lineage specification.

The craniofacial anomalies seen in individuals with Snijders Blok-Campeau syndrome suggest that CHD3 is essential for the specification and/or differentiation of CNCCs. In this study, we therefore sought to establish the role of CHD3 in craniofacial development using human iPSC models with either heterozygous or homozygous frameshift variants that result in loss of expression of the allele/gene. Importantly, established protocols are available to differentiate iPSCs into migratory CNCCs.^40–42^ With this approach we found that CHD3 is required to allow response to BMP during the specification of the CNCCs. Namely, CHD3 regulates accessibility at enhancers bound by BMP responsive transcription factors, including DLX and MSX families, as well as the expression of these factors. In the absence of CHD3, BMP response is not effective, and this leads to a Wnt/BMP imbalance. Thus, CNCC specification fails, replaced by mesodermal identity, which can be partially rescued by titrating Wnt levels.

## RESULTS

### CHD3 is not required for the pluripotent identity of the iPSCs

To investigate the role of CHD3 in CNCC specification, we used heterozygous and homozygous *CHD3* knock-out IPSC lines (and isogenic controls) generated in a companion study^43^ by means of CRISPR/Cas9 gene-editing in the BIONi010-A iPSC line (Fig. 1A). Specifically, for this study, we used two different homozygous clones (hereafter *CHD3*-KO clones 1 and 2) in which Cas9 independently targeted the third exon of the *CHD3* gene, producing a 1-base deletion (c.298delG), which generated a premature stop codon downstream (Fig. 1B).^43^ Moreover, we used two heterozygous clones (hereafter *CHD3*-HET-KO clones 1 and 2) that were generated using c.298insA and c.298insT respectively.^43^

**Figure 1.**
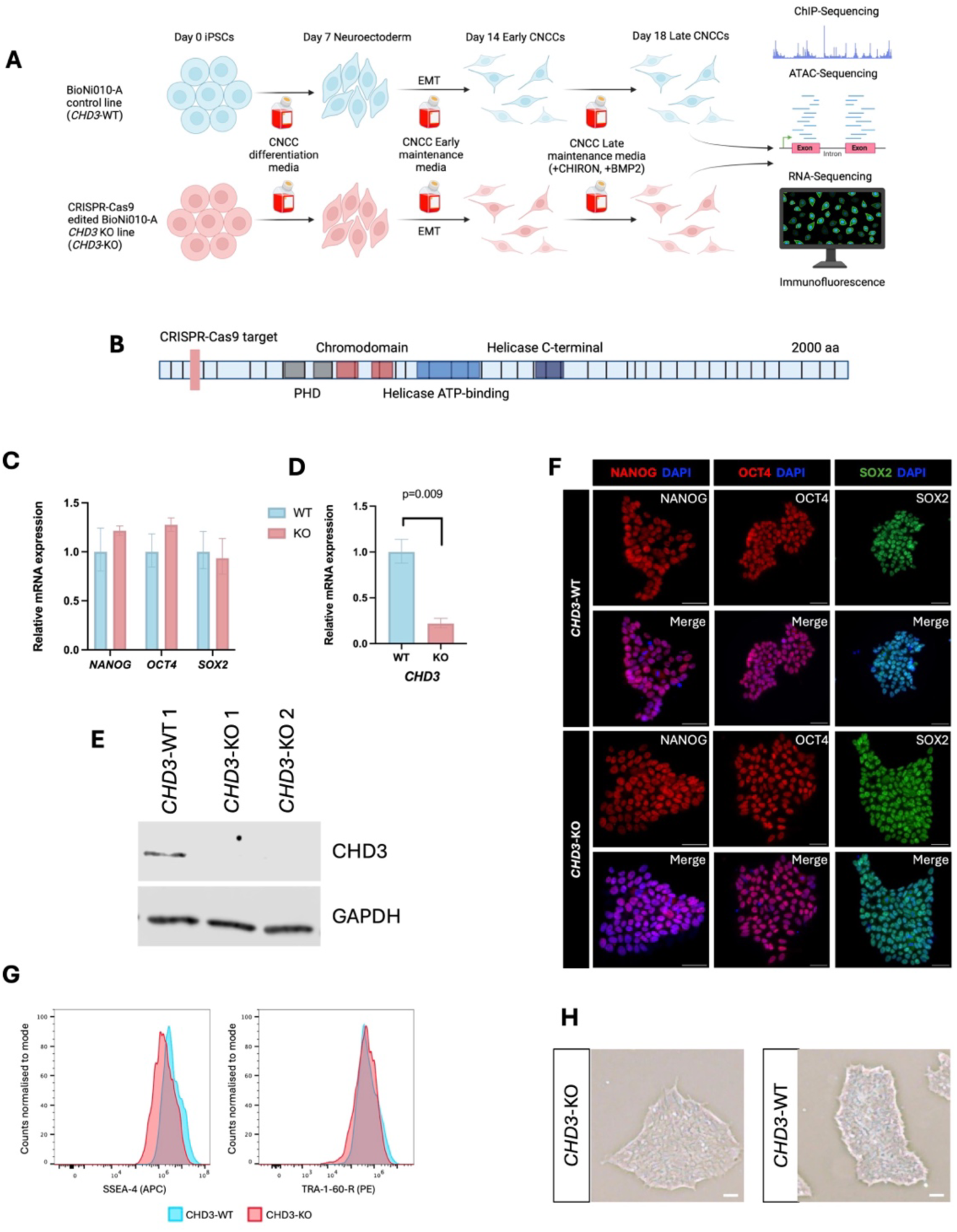
*CHD3*-KO lines are pluripotent. (A) A graphical illustration of the experimental pipeline. Made with biorender.com. (B) A schematic of the CHD3 gene indicating the site targeted by CRISPR-Cas9. Human CHD3 isoform 1, NM_001005273.2, 2000 aa, 40 exons (canonical). Made with biorender.com. (C and D) RT-qPCR quantifying the relative expression levels of (C) pluripotency markers and (D) *CHD3* in *CHD3*-WT and *CHD3*-KO iPSCs. Differences between lines were assessed using unpaired student’s t-test with significant p-values displayed. (E) Western blot for CHD3 in *CHD3*-WT clone 1 (*CHD3*-WT 1) and *CHD3*-KO clone 1 (*CHD3*-KO 1) and clone 2 (*CHD3*-KO 2) iPSCs. GAPDH is used as a loading control. (F) Immunofluorescence for key pluripotency markers in *CHD3*-WT and *CHD3*-KO iPSCs. Scale bar: 50 μm. (G) Flow cytometry for pluripotency surface markers in *CHD3*-WT and *CHD3*-KO iPSCs. (H) Example iPSC colonies from *CHD3*-KO and *CHD3*-WT lines. Scale bar: 50 μm.

In the original paper in which these CRISPR lines were generated ^43^, it was already established that neither the *CHD3*-KO nor the *CHD3*-HET-KO affect the pluripotency of the iPSCs. We further corroborated this finding in the present study with additional experiments. Specifically, RT-qPCR revealed no significant difference in the expression of the main pluripotency markers (*NANOG, OCT4* and *SOX2*) when comparing the *CHD3*-KO and *CHD3*-WT iPSCs (Fig. 1C). As expected, expression of *CHD3* was significantly lower in *CHD3*-KO iPSCs compared to *CHD3*-WT (Fig. 1D). Furthermore, loss of CHD3 at the protein level in the *CHD3*-KO iPSCs was also observed, confirming the success of the CRISPR knockout (Fig. 1E).^43^ RNA-seq performed at the iPSC stage found only 25 genes to be differentially expressed between *CHD3*-WT and *CHD3*-HET-KO iPSCs, and 62 genes to be differentially expressed between *CHD3*-WT and *CHD3*-KO iPSCs (FDR <5%; logFC > 1.5 or < -1.5). None of the known pluripotency factors were differentially expressed (Supplementary File S1). Immunofluorescence and flow cytometry analysis of several key pluripotency markers confirmed that protein levels of pluripotency factors are not affected by CHD3 loss (Fig. 1F, G). Overall, these data suggest that CHD3 loss has a modest impact on the iPSC transcriptome, and it does not affect the pluripotent gene and protein network. Consistent with this, *CHD3*-KO iPSCs exhibited regular morphology, forming tightly packed colonies with well-defined edges (Fig. 1H).

To further corroborate the pluripotent state of the *CHD3*-KO iPSCs, we performed trilineage differentiation and found that *CHD3*-KO iPSCs were able to successfully differentiate into all the three germ layers (Supplementary Figure S1).

In summary, these data confirm that the CRISPR knockout of *CHD3* was successful and that loss of CHD3 has no impact on iPSC pluripotent identity.

### Loss of CHD3 impairs CNCC specification

Next, we investigated whether *CHD*3-KO affects CNCC specification. To achieve this, we leveraged an established protocol^40,44^, which has been previously adapted by our lab.^41,42^ With this protocol, fully specified, migratory CNCCs are generated in 18 days.

We first tested the CNCC specification protocol on the *CHD3*-WT clones. At the endpoint of the differentiation (day-18), the cells expressed genes typical of CNCC identity (e.g. *SOX9, TFAP2A, TWIST1, NR2F1, SNAI1/2*; Fig. 2A), along with markers of EMT and mesenchymal state (e.g. *VIM, ZEB2, SNAI1/2, CDH2*; Fig. 2A). Conversely, epithelial and pluripotency markers were downregulated (e.g. *CDH1, POU5F1, NANOG, SOX2*; Fig. 2A). Overall, these data confirmed that both clones of *CHD3*-WT iPSCs were able to successfully differentiate into migratory CNCCs.

**Figure 2.**
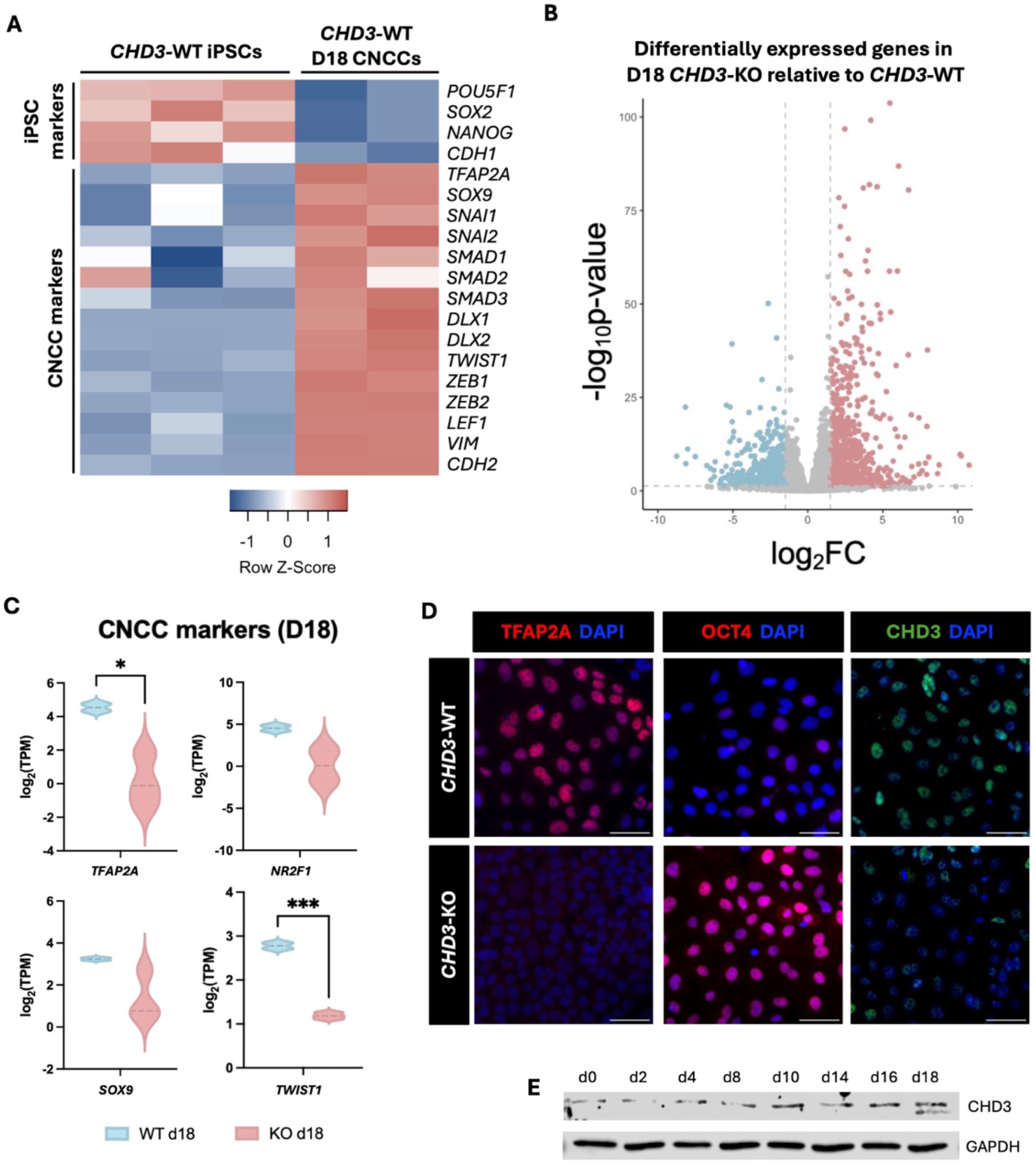
Loss of CHD3 impairs CNCC specification. (A) Heatmap displaying expression of key pluripotency markers and CNCC markers in *CHD3*-WT iPSCs and *CHD3-*WT Day-18 CNCCs. (B) Volcano plot of differentially expressed genes in *CHD3*-KO relative to *CHD3*-WT in Day-18 CNCCs. Blue dots represent downregulated genes with p-adj < 0.05 and log_2_FoldChange < -1.5. Red dots represent upregulated genes with p-adj < 0.05 and log_2_FoldChange > 1.5. (C) Violin plots produced from RNA-seq data generated in D18 *CHD3*-WT and *CHD3*-KO CNCCs. Log_2_(TPM) for the key CNCC markers (*TFAP2A, NR2F1, SOX9* and *TWIST1)* are displayed. P-values were obtained using a two-tailed unpaired t-test. *=p<0.05, **=p<0.01, ***=p<0.001. (D) Immunofluorescence for CNCC marker TFAP2A, pluripotency marker OCT4, and CHD3 in *CHD3*-WT and *CHD3-*KO D18 CNCCs. Scale bar: 50 μm. (E) Time-course western blot for CHD3 in *CHD3*-WT during the course of differentiation from iPSCs to CNCCs. GAPDH was used as a loading control.

We went on to investigate whether CHD3 loss had an impact on CNCC specification. To this end, we differentiated the *CHD3*-WT, *CHD3*-HET-KO, and *CHD3*-KO clones into CNCCs and conducted RNA-seq, paired with immunofluorescence for CNCC and iPSC markers. The cells were collected at the endpoint of the protocol (day-18). Comparing the two *CHD3*-WT clones with the *CHD3*-HET-KO counterparts, we found that heterozygous loss of *CHD3* did not have a major impact on CNCC specification. In fact, only 36 genes were differentially expressed in *CHD3*-HET-KO CNCCs relative to the *CHD3*-WT lines (FDR <5%; logFC > 1.5 or < -1.5; Supplementary Fig. S2; Supplementary File S2). This could potentially be due to compensation from the wild-type allele.

In stark contrast to the subtle effects of a heterozygous loss of *CHD3*, 1,516 genes were differentially expressed when comparing *CHD3*-KO and *CHD3*-WT CNCCs (FDR <5%; logFC > 1.5 or < -1.5; Fig. 2B). Of these, 862 genes were upregulated in the *CHD3*-KO CNCCs, while 654 were downregulated. Many CNCC markers were downregulated in *CHD3*-KO, including *TFAP2A*, *TWIST1*, *SOX9* and *NR2F1* (Fig. 2C). On the other hand, pluripotency genes such as *POU5F1* (OCT4) and *NANOG* were upregulated (Fig. 2C; Supplementary File S3). TFAP2A downregulation and OCT4 upregulation in *CHD3*-KO CNCCs were detected also at the protein level (Fig. 2D). Moreover, epithelial genes (e.g. *EPCAM* and *CDH1*) were upregulated in the *CHD3*-KO cells, while mesenchymal genes (*VIM*, *CDH2*) were downregulated. These findings suggest that the *CHD3*-KO cells failed to induce the cadherin switch (from high CDH1/low CDH2 to low CDH1/high CDH2) which usually correlates with successful EMT in CNCCs. Finally, we noted that the two other NuRD paralogs *CHD4* and *CHD5* were not upregulated in *CHD3*-KO CNCCs, potentially excluding compensatory mechanisms between NuRD subunits.

Together, these data suggest that CHD3 has an important role in CNCC specification. This role is reflected by the gradual upregulation of this protein, from relatively low levels in iPSCs and during neuroectoderm formation (∼days 1–8; Fig. 2E), to higher levels in pre-migratory and migratory CNCCs (days 10–18; Fig. 2E). The CHD3 protein upregulation reflects a progressive upregulation in *CHD3* gene expression as suggested by RNA-seq performed in iPSCs (*CHD3* median TPM = 3.4), early CNCCs (day-14, median TPM = 11.5) and late, fully specified CNCCs (median TPM = 18.8). Interestingly, a slightly shorter isoform of *CHD3* (ENST00000358181.8), lacking the original exons 1 (replaced with an alternative exon 1) and 33 is expressed at relatively low levels in fully specified CNCCs (day-18 median TPM = 3.3), while it is not expressed at the previous time points. The expression of the shorter isoform at the day-18 is observable also at the protein level (Fig. 2E).

### *CHD3*-KO cells undergo mesodermal fate

To shed light on the function of CHD3 in CNCC specification, we performed Gene Ontology (GO) analysis on the 1,516 genes differentially expressed in *CHD3*-KO CNCCs on day-18. GO terms downregulated in the *CHD3*-KO CNCCs were mainly associated with development, morphogenesis, patterning, and cell motility (Fig. 3A). Conversely, the upregulated genes were enriched for cell-cell junction and ion channel terms, potentially explaining the impaired EMT process and the persistent epithelial state of the *CHD3*-KO cells (Fig. 3B). Importantly, the *CHD3*-KO cells also exhibited upregulation of genes typically expressed in the primitive streak and in the early pre-migratory mesoderm, such as *EOMES, TBXT, TBX3, TBX6*, and *MIXL1*, paired with downregulation of BMP responsive transcription factors, including several paralogs of the DLX and MSX families (Figs. 3C–E).^30,45,46^

**Figure 3.**
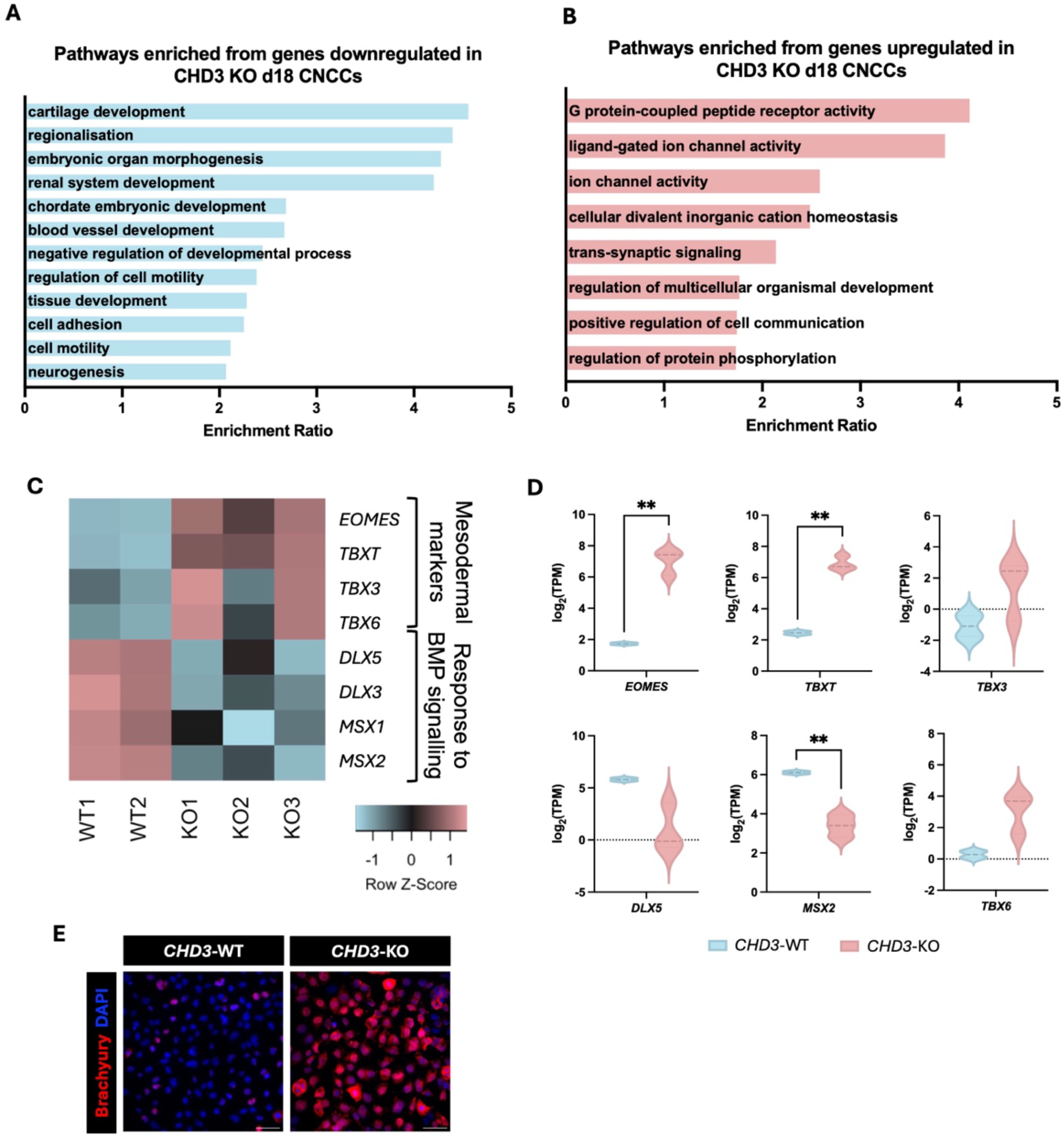
*CHD3*-KO CNCCs display aberrant expression of early mesodermal markers. (A and B) GO term analysis of (A) significantly downregulated and (B) significantly upregulated genes in D18 *CHD3*-KO CNCCs as determined using WebGestalt pathway analysis. (C) Heatmap displaying expression of mesodermal markers and genes involved in BMP signalling in *CHD3*-WT (WT) and *CHD3*-KO (KO) D18 CNCCs. (D) Violin plots produced from RNA-seq data obtained from D18 *CHD3*-WT and *CHD3*-KO CNCCs. Log_2_(TPM) for key mesodermal markers (*EOMES, TBXT, TBX3* and *TBX6)* and BMP responsive transcription factors (*DLX5* and MSX2) are displayed. P-values were obtained using a two-tailed unpaired t-test. **=p<0.01. (E) Immunofluorescence for the mesodermal marker brachyury in *CHD3*-WT and *CHD3-*KO D18 CNCCs. Scale bar: 50 μm.

### Chromatin accessibility is impaired at BMP responsive enhancers

Our results so far indicate that the *CHD3*-KO cells display an aberrant early mesoderm signature, while failing to upregulate genes associated with BMP signalling, developmental patterning and CNCC identity. Next, we sought to elucidate the underlying molecular mechanism.

CHD3 is part of the NuRD chromatin remodeling complex, therefore we reasoned that the observed phenotypes could be caused by dysregulation of chromatin accessibility. To test this, we performed ATAC-seq on *CHD3*-WT and *CHD3*-KO clones at the endpoint of the CNCC specification protocol (day-18). We identified 23,121 ATAC-seq peaks which were conserved between all *CHD3*-WT and *CHD3*-KO clones (FDR <5%; Fig. 4A–C). On the other hand, 17,543 peaks were found exclusively in the *CHD3*-WT CNCCs but not in *CHD3*-KO counterparts (hereafter, *CHD3*-WT specific peaks), while 3,350 peaks were found to be exclusive to the *CHD3*-KO CNCCs (hereafter *CHD3*-KO specific peaks; Fig. 4A–C).

**Figure 4.**
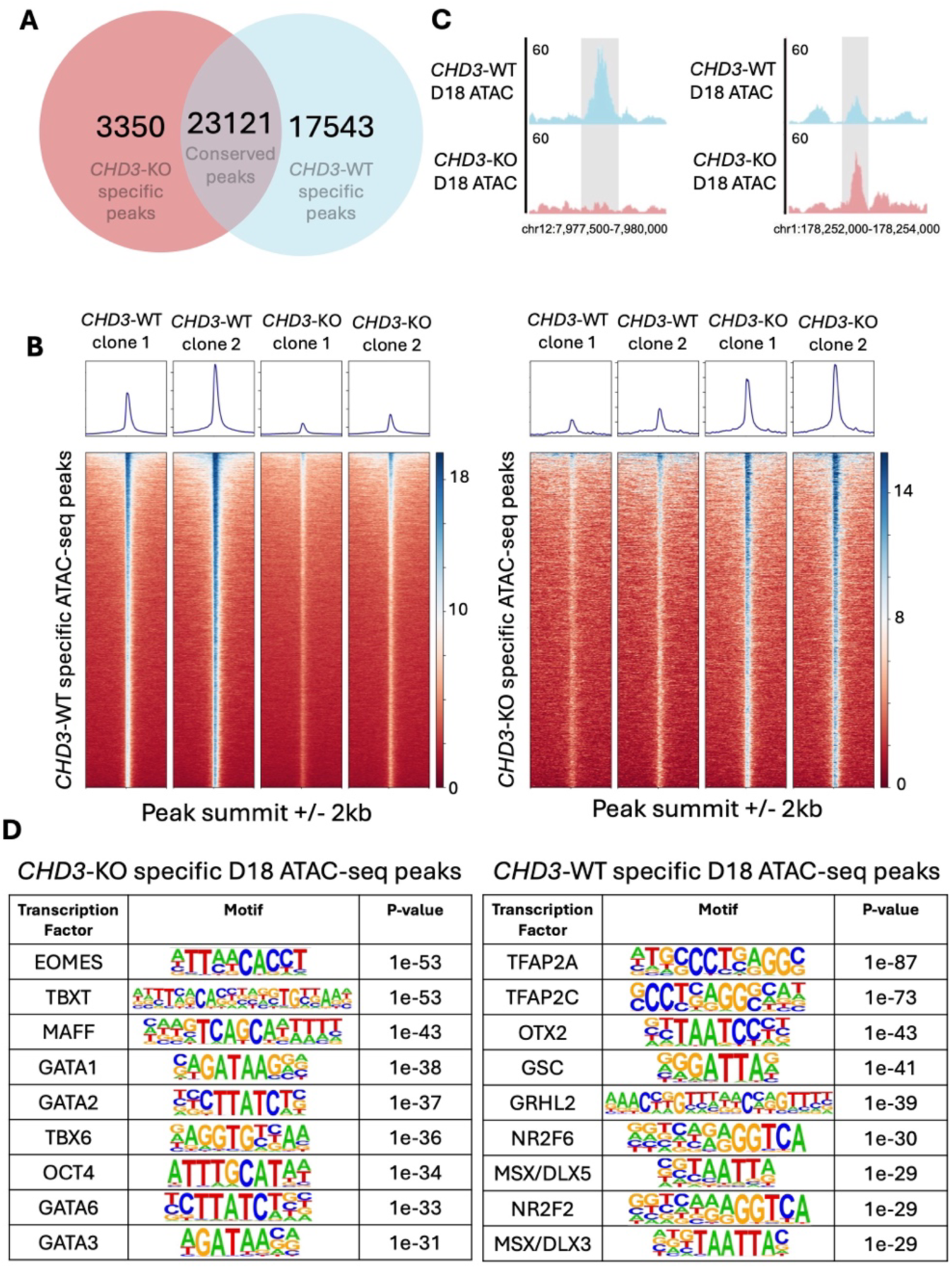
CHD3 loss alters chromatin accessibility in D18 CNCCs. (A) Venn diagram showing the number of *CHD3*-KO specific ATAC-seq peaks, *CHD3*-WT specific ATAC-seq peaks and conserved (peaks present in both *CHD3*-WT and *CHD3*-KO) ATAC-seq peaks in Day-18 CNCCs. (B) Heatmaps showing ATAC-seq peaks present in individual *CHD3*-WT and *CHD3*-KO replicates which are *CHD3*-WT or *CHD3*-KO specific. (C) Example of *CHD3*-WT specific and *CHD3*-KO specific ATAC-seq peaks visualised in UCSC genome browser. (D) Tables of motifs enriched in either *CHD3*-KO specific Day-18 ATAC-seq peaks or *CHD3*-WT specific D18 ATAC-seq peaks identified using HOMER.

Overall, 90% of the *CHD3*-WT specific and 97% of the *CHD3*-KO specific ATAC-seq peaks were distal from the nearest Transcription Start Site (TSS; i.e. distance > 1 Kb), suggesting that most of these regions are putative enhancers. Hence, these results indicate that loss of CHD3 had a significant impact on chromatin accessibility at enhancers.

Notably, 382 of the 654 genes downregulated in *CHD3*-KO cells (58%) were also the nearest gene to an enhancer that lost accessibility in these cells (i.e. *CHD3*-WT specific ATAC-seq peak), while 226 of the 862 upregulated genes (26%) were also the closest gene to an enhancer that gained accessibility. This pattern suggests that CHD3 regulates a significant number of genes through direct regulation of chromatin accessibility at their enhancers, acting both as activator and repressor, consistent with recent studies that suggested that the NuRD complex can both activate and repress gene expression in a context specific manner.^8–10^ Motif analysis of the enhancers aberrantly active in *CHD3*-KO cells showed enrichment for binding motifs of several primitive streak and early mesoderm transcription factors, including EOMES, TBXT (BRACHYURY), TBX3/6, and GATA (Fig. 4D).^47–50^ On the other hand, the enhancers aberrantly repressed in *CHD3*-KO cells were enriched for motifs of transcription factors implicated in CNCC specification (TFAP2A/C, NR2F2/6), BMP response, and patterning (DLX/MSX families and OTX2) ^28–30,45,51^.

In summary, both the RNA-seq and the ATAC-seq consistently demonstrated that the CHD3-deficient cells induced an aberrant mesodermal program, while failing to activate CNCC specification possibly due to impaired response to BMP signalling.

### CHD3 primes BMP response in the developing CNCCs

Our experiments conducted in terminally specified CNCCs (day-18) indicated that BMP signalling response might be dysregulated in *CHD3*-KO conditions and that the cells acquire an unexpected mesodermal signature. It has previously been established that a combination of Wnt and FGF promotes the differentiation of iPSCs into mesodermal lineages.^52–56^ Consistent with this, in our protocol, the iPSCs are initially treated with FGF alone, while a Wnt agonist (CHIR99021, hereafter CHIRON) is added together with BMP2 at day-14 of differentiation (i.e. after 2-3 passages from the emergence of the first CNCCs). As aforementioned, BMP signalling is crucial to enable CNCC specification and differentiation.^26,30,35–39,57^ In particular, a fine-tuned balance between Wnt and BMP pathways is essential for proper CNCC specification.^53,58,59^

Based on this premise, we surmised that the mesodermal signature observed in the *CHD3*-KO cells might be caused by exposure to the Wnt agonist paired with the inability of the cells to properly respond to BMP2 stimuli, leading to a Wnt/BMP imbalance. Consistent with this, RNA-seq performed at day-14 of the CNCC specification protocol (i.e. right before exposure to CHIRON) revealed that the mesodermal markers are not yet expressed in the *CHD3*-KO cells before Wnt exposure (Fig. 5A). This finding supports the hypothesis that the primitive streak and mesodermal genes are eventually induced by the Wnt pulse.

**Figure 5.**
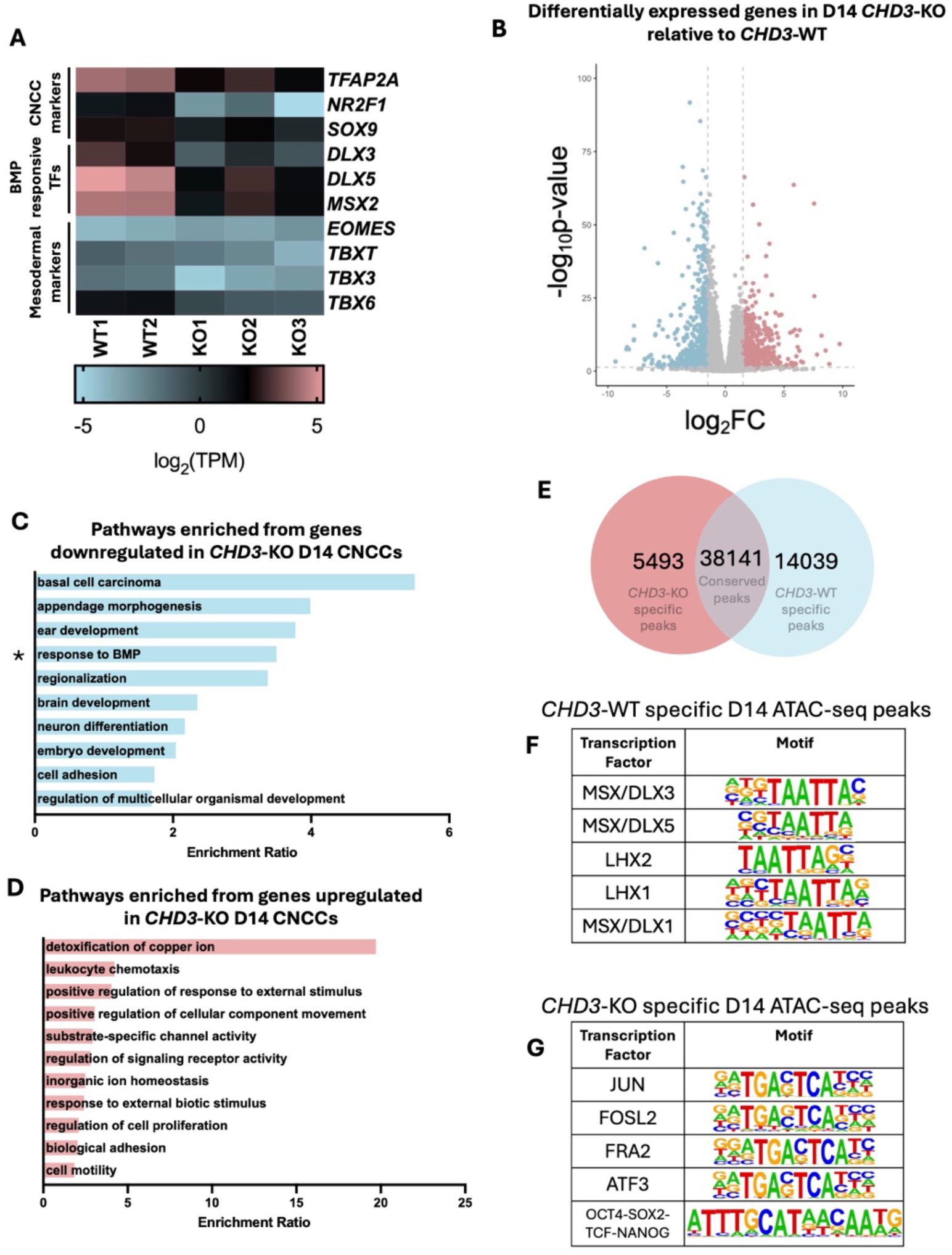
The effects of CHD3 loss on gene expression and chromatin accessibility in D14 CNCCs. (A) Heatmap displaying expression of CNCC markers (*TFAP2A, NR2F1,* SOX9), genes involved in BMP signalling (*DLX3, DLX5, MSX2*) and mesodermal specification (*EOMES, TBXT, TBX3, TBX6*) in *CHD3*-WT (WT) and *CHD3*-KO (KO) D14 CNCCs. (B) Volcano plot of differentially expressed genes in *CHD3*-KO relative to *CHD3*-WT in Day-14 CNCCs. Blue dots represent downregulated genes with p-adj < 0.05 and log_2_FoldChange < -1.5. Red dots represent upregulated genes with p-adj < 0.05 and log_2_FoldChange > 1.5. (C and D) GO term analysis of (C) significantly downregulated and (D) significantly upregulated genes in D14 *CHD3*-KO CNCCs as determined using WebGestalt pathway analysis. (E) Venn diagram showing the number of *CHD3*-KO specific ATAC-seq peaks, *CHD3*-WT specific ATAC-seq peaks and conserved (peaks present in both *CHD3*-WT and *CHD3*-KO) ATAC-seq peaks in Day-14 CNCCs. (F and G) Tables of motifs enriched in either (F) *CHD3*-KO specific Day-14 ATAC-seq peaks or (G) *CHD3*-WT specific Day-14 ATAC-seq peaks identified using HOMER.

Genome-wide analysis of the transcriptome at the day-14 timepoint identified 1,411 differentially expressed genes (FDR <5% and fold change >1.5 or <-1.5; Supplementary File S4), with 569 being upregulated in the *CHD3*-KO and 842 downregulated (Fig. 5B). GO term analysis of the 842 downregulated genes revealed significant enrichment for BMP signalling response, morphogenesis, and patterning (Fig. 5C, D).

We next performed ATAC-seq at the same timepoint (day-14). Similar to day-18, at day-14 thousands of chromatin regions were either aberrantly open or aberrantly closed in the *CHD3*-KO cells (14,039 and 5,493 regions respectively, FDR < 5%; Fig. 5E). Strikingly, the homeobox motif associated to the DLX/MSX BMP-responsive factors was the only motif enriched in the 5,493 aberrantly closed regions at day-14 (i.e. day-14 *CHD3*-KO specific peaks; Figs. 5F, G; regions whose accessibility was not affected by CHD3 loss were used as control for the differential motif analysis).

Overall, the experiments conducted at day-14 suggest that CHD3 may have the role of priming the developing CNCCs to respond to BMP by opening the chromatin at the BMP responsive enhancers, facilitating the binding of homeodomain factors.

### CHD3 binds the BMP responsive enhancers

Next, we set out to investigate if CHD3 binds directly at the enhancers that either lose or gain chromatin accessibility in the *CHD3*-KO cells, both at day-14 (before Wnt and BMP exposure) and day-18 (after Wnt and BMP exposure). We thus performed ChIP-seq for CHD3 at these two time points in *CHD3*-WT cells and detected CHD3 binding at nearly all the *CHD3*-WT-specific and *CHD3*-KO-specific ATAC-seq peaks at both time points (Fig. 6 A–E).

**Figure 6.**
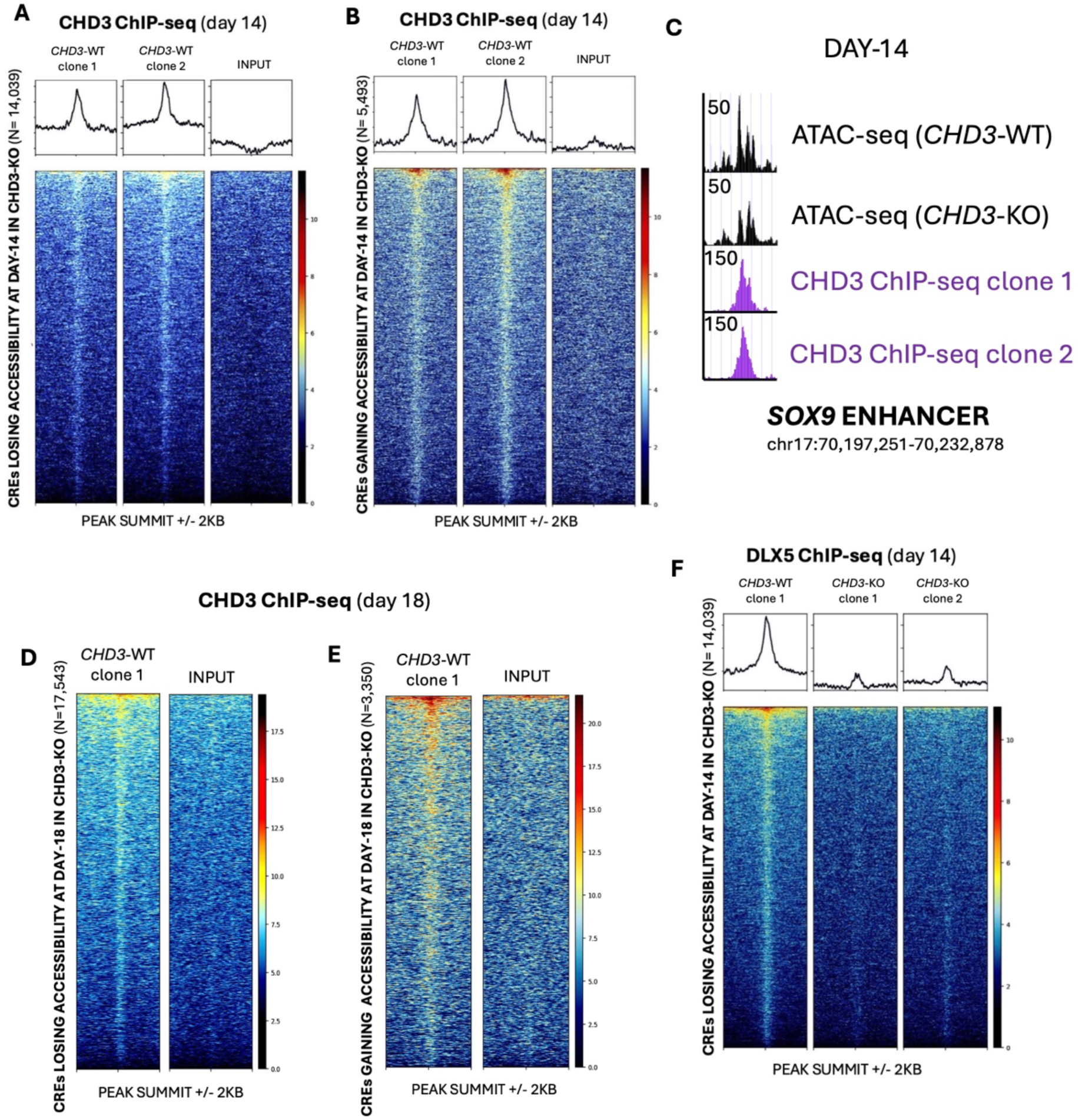
CHD3 binding in CNCCs correlates with regions where chromatin accessibility is altered upon CHD3 loss. (A and B) Heatmap of CHD3 binding at cis-regulatory elements (CREs) which either (A) lose accessibility or (B) gain accessibility in Day-14 *CHD3*-KO CNCCs. (C) An example of an enhancer bound by CHD3 in Day-14 CNCCs which loses accessibility in *CHD3*-KO. (D and E) Heatmap of CHD3 binding at cis-regulatory elements (CREs) which either (D) lose accessibility or (E) gain accessibility in D18 *CHD3*-KO CNCCs. (F) Heatmap of DLX5 binding at cis-regulatory elements (CREs) which lose accessibility in Day-14 *CHD3*-KO CNCCs.

Given that most of the sites that are aberrantly closed at both day-14 and day-18 are enriched for the binding motif of the DLX/MSX factors, we performed ChIP-seq for DLX5 at day-14 in *CHD3*-WT and *CHD3-*KO cells. We specifically selected DLX5 because it is highly expressed in our cell model at this timepoint and it has been previously associated with BMP response, CNCC specification and craniofacial development.^45,60–62^ This experiment confirmed that DLX5 normally binds at these sites at day-14 and that loss of CHD3 results in depletion of DLX5 binding from these sites (Fig. 6F), possibly because of the significantly reduced chromatin accessibility, paired with downregulation of the *DLX5* gene.

In summary, our data indicate that CHD3 directly binds at important BMP responsive enhancers, and that loss of CHD3 binding from these regulatory elements attenuates their accessibility, ultimately affecting the ability of crucial transcription factors to bind. This leads to impairment in BMP response, which triggers an imbalance between signalling pathways.

### Titration of Wnt levels attenuates the aberrant mesodermal signature

Finally, we investigated whether attenuating Wnt signalling could rescue the aberrant phenotypes observed in *CHD3*-KO conditions. To achieve this, we differentiated *CHD3*-WT and *CHD3*-KO cells into CNCCs using different CHIRON concentrations: 3, 2 and 1 μM. At 3 μM (original protocol), the *CHD3*-KO cells displayed high expression of mesodermal markers both at gene and protein levels, paired with significantly reduced expression of CNCC markers (Fig. 7 A, B). Notably, decreasing CHIRON (2 μM and 1 μM respectively) was sufficient to significantly lower the activation of the mesodermal genes (e.g. *EOMES* and TBX3; Fig. 7 A, B) but it was ineffective in restoring the expression of the CNCC markers (Fig. 7 A, B).

**Figure 7.**
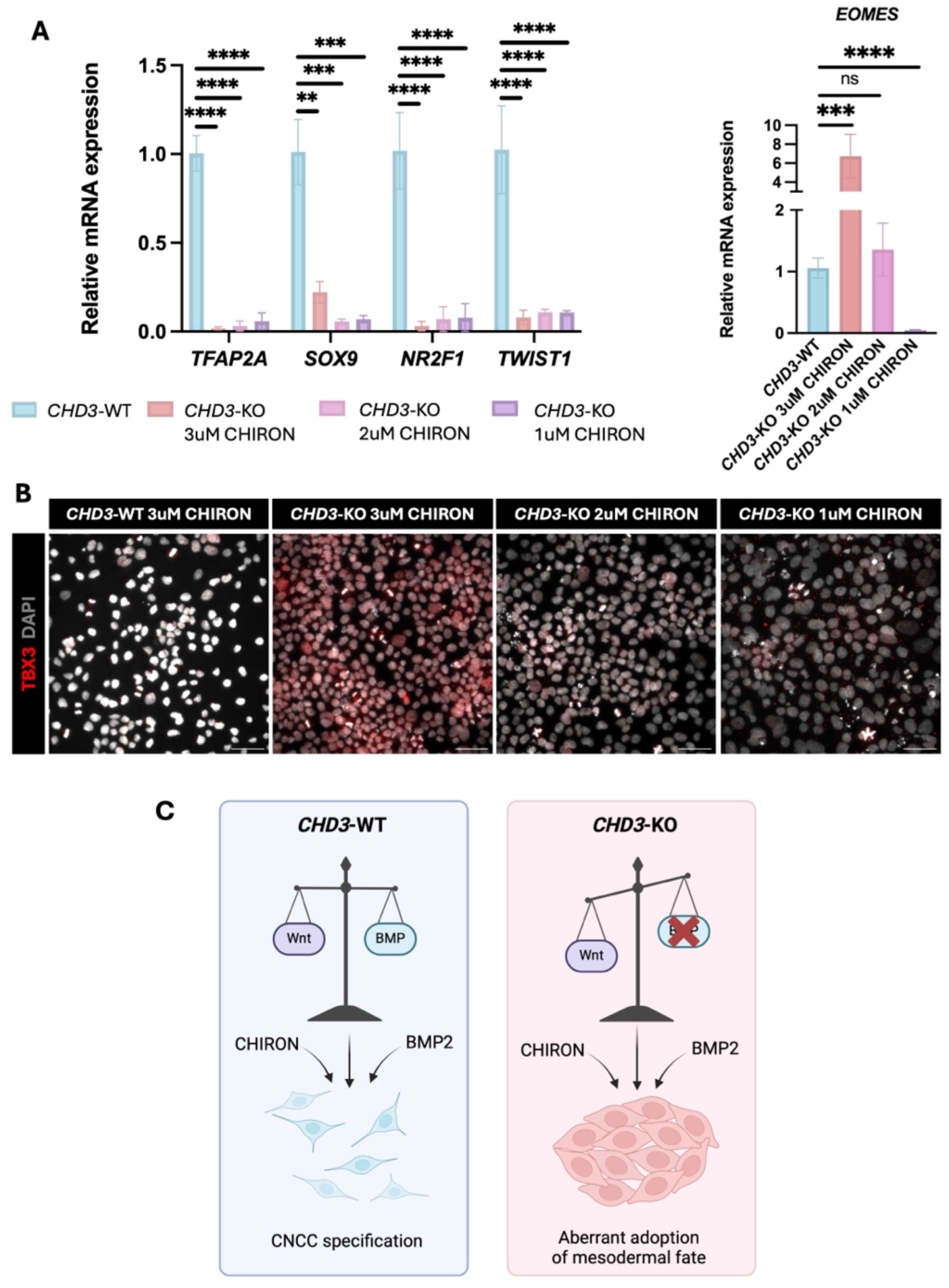
*CHD3-*KO aberrant mesodermal phenotype is rescued by attenuated Wnt signalling. (A) RT-qPCR assessing the relative expression levels of CNCC markers (*TFAP2A, SOX9, NR2F1* and *TWIST*) and mesodermal marker (*EOMES*) between *CHD3*-WT and *CHD3*-KO Day-18 CNCCs provided with either the standard 3μM or a reduced concentration (2μM or 1μM) of CHIRON. Differences between conditions were assessed using unpaired student’s t-test. **=p<0.01, ***=p<0.001, ****=p<0.0001, ns=not significant. (B) Immunofluorescence for the mesodermal marker TBX3 in *CHD3*-WT and *CHD3-*KO Day-18 CNCCs provided with either the standard 3μM or a reduced concentration (2μM or 1μM) of CHIRON. Scale bar: 50 μm. (C) Graphical summary of the effect of CHD3 loss on CNCC specification. Loss of CHD3 in *CH3-*KO disrupts the BMP signalling response leading to an imbalance between Wnt and BMP signalling, ultimately resulting in adoption of an aberrant mesodermal fate. Made with biorender.com.

Next, we tried to rescue the expression of the CNCC markers in *CHD3*-KO cells by simultaneously decreasing Wnt (1 μM of CHIRON) and increasing BMP (BMP2 concentration raised from 50 to 150 pg/ml). However, even subjecting the cells to three times more BMP2 was not effective in restoring the expression of the CNCC genes (Supplementary Fig. S3), suggesting that the response to BMP is permanently impaired by the loss of CHD3.

Overall, these data support a model in which the aberrant primitive streak/early mesoderm signature observed in *CHD3*-KO cells is caused by a Wnt/BMP imbalance (Fig. 7C) which can be overcome by attenuating Wnt levels. On the other hand, the impaired CNCC specification is the result of ineffective BMP response, which cannot be rescued by simply increasing the amount of BMP ligand.

## DISCUSSION

CHD3, a chromodomain helicase DNA-binding protein, is a core component of the NuRD complex, which modulates chromatin structure to regulate transcription.^63,64^ The NuRD complex is essential for developmental processes as it coordinates histone deacetylation and nucleosome remodeling to ensure precise gene expression patterns.^7,65^ Heterozygous pathogenic variants in *CHD3* cause Snijders Blok-Campeau syndrome, a rare autosomal dominant neurodevelopmental syndrome characterized by a complex array of phnotypes that vary in severity between different affected individuals, including variable degrees of intellectual disability, impaired speech, macrocephaly, and distinct craniofacial features.^19,20^ Many of the known pathogenic variants occur in the ATPase domain, which likely compromises chromatin remodeling and disrupts developmental gene expression programs, and loss-of-function alleles have also been reported.^19,20^ The observations of distinctive craniofacial dysmorphisms in individuals with Snijders Blok-Campeau syndrome suggest that CHD3 plays a pivotal role in cranial neural crest cell (CNCC) development.^19,20^ Although CHD3 has been implicated in neuronal migration and synapsis formation during brain development^18,43^, its potential roles in CNCC specification and differentiation had never been investigated prior to the present study.

In this study, we demonstrate that CHD3 is indispensable for the proper specification of human CNCCs. Using CRISPR-Cas9-based knockout models, we showed that iPSCs with homozygous *CHD3* knockout retain pluripotent identity but exhibit severe defects in CNCC specification. Transcriptomic analysis revealed significant downregulation of CNCC markers such as *TFAP2A*, *SOX9*, and *TWIST1* in *CHD3*-KO cells, paired with upregulation of mesodermal markers like *TBXT*, *EOMES* and several others. These findings were corroborated by chromatin accessibility assays, which showed that a complete lack of CHD3 protein leads to the loss of open chromatin at enhancers bound by BMP-responsive transcription factors, such as the DLX and MSX families, essential for CNCC specification.^45,46^ Furthermore, our data indicate that *CHD3*-KO cells fail to execute the epithelial-to-mesenchymal transition (EMT), as evidenced by the persistent expression of epithelial markers and failure to upregulate mesenchymal markers. These results suggest that CHD3 might be essential also for chromatin remodeling at loci regulating EMT and mesoderm-ectoderm fate decisions.^51,66^

Our findings provide critical insights into the Wnt/BMP balance, a key regulatory axis during embryogenesis. BMP signalling is crucial for CNCC specification, acting through SMAD-dependent transcriptional programs to induce patterning genes such as *MSX1* and *DLX5.^29,30,35^* The specific role of BMP during the different stages of CNCC specification, migration and differentiation has been debated. Studies in *Xenopus* and zebrafish have suggested that BMP gradients are required to produce a specific level of BMP signalling which is permissive for neural crest formation.^67–69^ However, there also other studies in *Xenopus*, zebrafish, and chicks which suggest that cranial neural crest induction is dependent on an initial inhibition of BMP signalling, followed by BMP activation.^70–72^ In particular, BMP4 and BMP7 have been implicated in early cranial neural crest specification.^26^ BMP signalling continues to play a key role in CNCC formation, enabling migration and later differentiation into derivative cells.^36^ Here, BMP2 appears to have an essential function in establishing migratory CNCCs^37,38^, while BMP2, BMP4 and BMP7 are critical in enabling the subsequent formation of craniofacial structures.^39^ In our study, *CHD3*-KO cells exhibited a marked reduction in BMP-responsive gene expression and chromatin accessibility, indicating that CHD3 is required for the transcriptional activation of BMP target genes.

Wnt signalling promotes mesodermal differentiation by stabilizing beta-catenin, which activates mesoderm-specific transcription factors such as *EOMES*, *TBXT* and *GATA3.*^48,52^ In *CHD3*-KO cells, elevated Wnt signalling led to the aberrant upregulation of mesodermal markers, suggesting that CHD3 modulates the interplay between Wnt and BMP to ensure proper CNCC fate determination. FGF signalling, which synergizes with Wnt to promote mesodermal differentiation, also appeared to contribute to the aberrant mesodermal fate of *CHD3*-KO cells.^58^ Notably, reducing Wnt levels attenuated the mesodermal signature, supporting the hypothesis that *CHD3*-deficient cells are unable to balance the Wnt signalling in the absence of BMP response. However, this experiment failed to restore the expression of the CNCC markers, emphasizing that CHD3 is indispensable for BMP signal transduction, independently from the relative contributions of the other signalling pathways.

This study establishes CHD3 as a pivotal chromatin remodeler that integrates BMP and Wnt signalling to regulate CNCC fate decisions. By modulating chromatin accessibility at BMP-responsive enhancers, CHD3 ensures the proper transcriptional activation of key developmental genes. Dysregulation of this balance, as seen in *CHD3*-KO cells, leads to anomalous mesodermal differentiation, providing a potential mechanistic explanation for the craniofacial defects observed in individuals with Snijders Blok-Campeau syndrome.

Importantly, the companion study where the CRISPR-iPSC lines were generated^43^ investigated the role of CHD3 in cortical development, and found that CHD3 is highly expressed in mature neurons, where it regulates synaptic development and function, suggesting that CHD3 may have different roles in different developmental processes.

Our work also suggests that the contributions of CHD3 to embryonic stem cell pluripotency are negligible, likely reflecting its relatively low expression levels in this cell type, where the paralog CHD4 has been demonstrated to be dominant and essential.^73–75^ Importantly, we did not find evidence of compensation to CHD3 loss by upregulation of other CHD paralogs such as CHD4 or CHD5, suggesting that the function of mediating BMP response might be exclusive of CHD3.

Future research should explore the interactions between CHD3-NuRD and other chromatin remodeling complexes, as well as its potential role in fine-tuning Wnt/FGF signaling during other steps of craniofacial development. In particular, given the importance of BMP in craniofacial osteogenesis, future studies should also explore the function of CHD3 in the formation of CNCC-derived craniofacial bones and cartilage. Additionally, *in vivo* studies using model organisms could further elucidate the developmental contexts in which CHD3 operates.

### Limitations of the study

This study was conducted using *CHD3* CRISPR-KO models and showed that *CHD3* heterozygous KO has no effect on CNCC specification, while *CHD3* homozygous KO has dramatic consequences on the same developmental process. However, it is important to highlight that a significant fraction of the individuals with Snijders Blok-Campeau syndrome present with heterozygous *CHD3* missense variants. We speculate that these heterozygous missense variants could have a dominant negative effect, which is recapitulated *in vitro* by complete (homozygous) loss of CHD3, as in our system. Therefore, the present study would ideally be complemented by future research that employs cells from affected individuals carrying the specific heterozygous missense variants in the *CHD3* gene, or isogenic lines engineered to carry those same variants.

Finally, the “TAATTA” sequence, whose chromatin accessibility in CNCCs is regulated by CHD3, is recognised as a binding motif by most homeodomain factors, and not just DLX5. It is part of the CNCC “coordinator motif”^40^ and consequently we cannot exclude that other homeodomain factors (e.g. ALX family^76^) could also be implicated in CHD3-mediated BMP response.

## RESOURCE AVAILABILITY

### Lead contact

Further information and requests for resources and reagents should be directed to and will be fulfilled by the lead contact, Marco Trizzino.

### Materials availability

Inquiries on CRISPR cell lines used in this study should be directed to Dr. Simon E. Fisher.

### Data and code availability

RNA-seq, ATAC-seq, and ChIP-seq data have been deposited in the Gene Expression Omnibus (GEO) under accession code GEO: GSE288669 and are publicly available as of the date of manuscript submission. Any additional information required to re-analyze the data reported in this paper is available from the lead contact upon request.

## ACKNOWLEDGMENTS

We thank Dr. Samantha Barnada (Thomas Jefferson University) and Prof. Brian Hendrich (University of Cambridge) for insightful discussions on the data. The authors thank the Genomic Facility at The Wistar Institute (Philadelphia, PA) for the Next Generation Sequencing. For this work, MT was funded by the G. Harold and Leila Y. Mathers Foundation and by BBSRC.

## AUTHOR CONTRIBUTIONS

MT, SEF and PC designed the project. ZHM performed most of the experiments. JdH and WC generated and performed validations of the CRISPR iPSC lines. MD and LD contributed to some of the experiments. ZHM and MT analyzed the data and wrote the manuscript. KL contributed to data analysis and offered critical support. All the authors read and approved the manuscript.

## DECLARATION OF INTERESTS

The authors declare no competing interests.

## METHODS

### Generation of the CRISPR iPSC lines

Heterozygous and homozygous *CHD3* knock-out IPSC lines were generated via CRISPR/Cas9 gene-editing in a companion paper.^43^ Specifically, for this study, we used two different homozygous clones (*CHD3*-KO) in which Cas9 targeted the third exon of the *CHD3* gene in the established BIONi010-A iPSC line, producing a 1-base deletion (c.298delG) which generated a premature stop codon downstream (Fig. 1B).^43^ Similarly, two heterozygous clones (*CHD3*-HET-KO) were instead generated using c.298insA and c.298insT respectively^43^, using the same BIONi010-A iPSC line.

### Culturing of human iPSCs and differentiation into CNCCs

*CHD3*-WT, *CHD3*-HET-KO and *CHD3*-KO lines were expanded through culturing of the iPSCs on geltrex (Thermo Fisher Scientific, A1413302) coated wells in mTesR plus medium (Stem Cell Technologies, 100-0276) containing 1% penicillin-streptomycin (Gibco, 15070063). The iPSCs were subsequently differentiated into CNCCs following the method developed by Prescott *et al.*^40^ Briefly, cells were cultured in CNCC differentiation media (table 1) for 6 days. On day 7-10, CNCC early maintenance media (table 1) was introduced and the cells were cultured in this up to day 14. On day 15, CNCC late maintenance media (table 1) was added, and cells were maintained in this up to day 18. Throughout differentiation, media was changed every other day and cells were passaged using StemPro accutase (Gibco, A1110501) each time 80% confluency was reached.

**Table 1.**
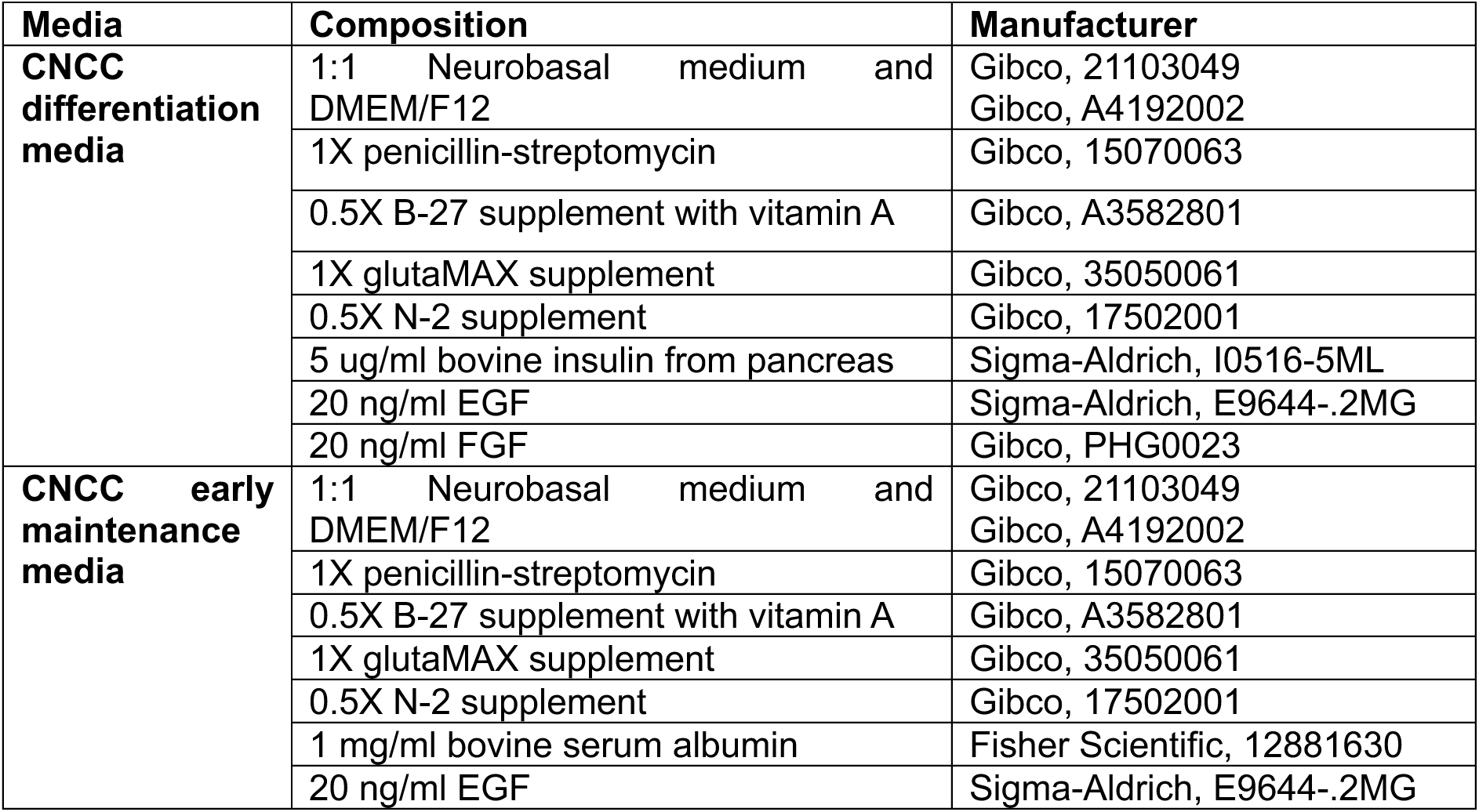
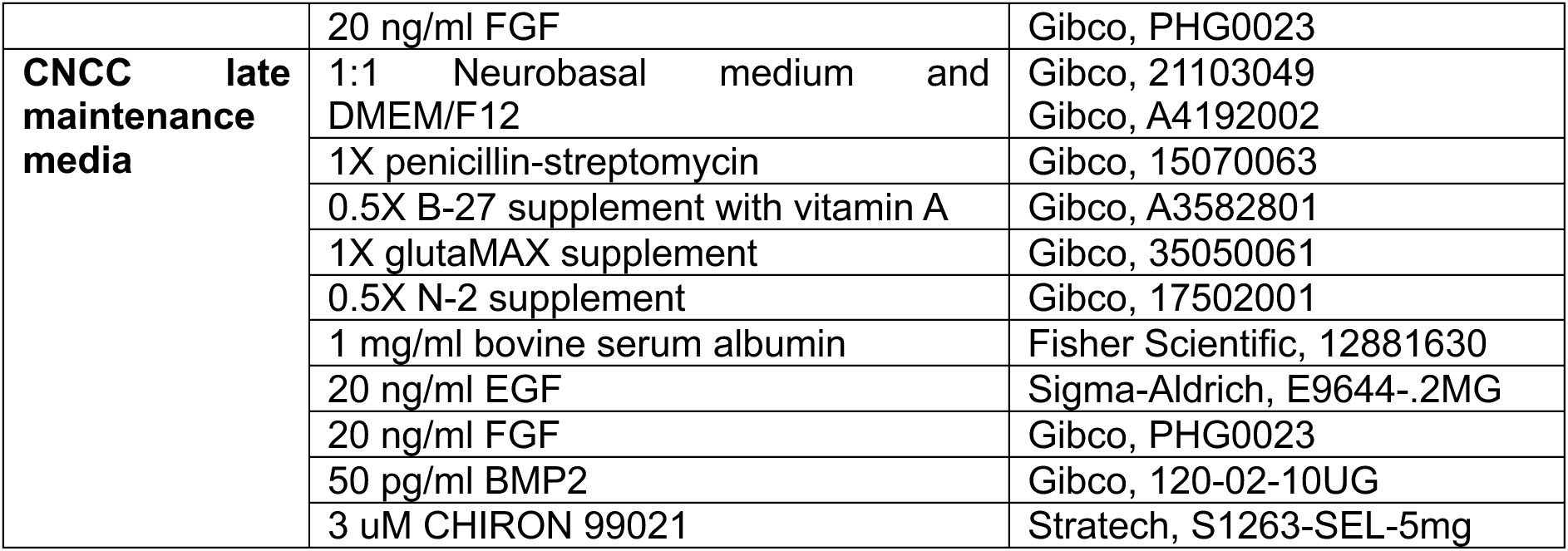
Media used for differentiation of iPSCs into CNCCs.

### Reverse transcription quantitative polymerase chain reaction (RT-qPCR)

Cells were pelleted and RNA was extracted using the Monarch Total RNA Miniprep Kit (NEB, T2010). 600ng of the extracted RNA was then converted to cDNA using the Thermo Scientific Maxima First Stand cDNA Synthesis Kit for RT-qPCR (Thermo Scientific, K1641). The qPCR reactions were prepared in a 96-well plate with each well containing 7.5ng cDNA, 5ul PowerUp SYBR Green Master Mix for qPCR (Applied Biosystems, A25778), 0.5uM each of forward and reverse primers (table 2) and 1.5ul of water for a total reaction volume of 10ul. The qPCR was carried out using a Bio-rad Connect qPCR machine with the following conditions: 3 minutes at 95°C, followed by 40 cycles of 10 seconds at 95°C, 20 seconds at 63°C and 30 seconds at 72°C, with a final melt curve of 65°C to 95°C for 5 minutes. Technical and biological replicates were carried out for each sample and 18S rRNA was used to normalise samples.

**Table 2.**
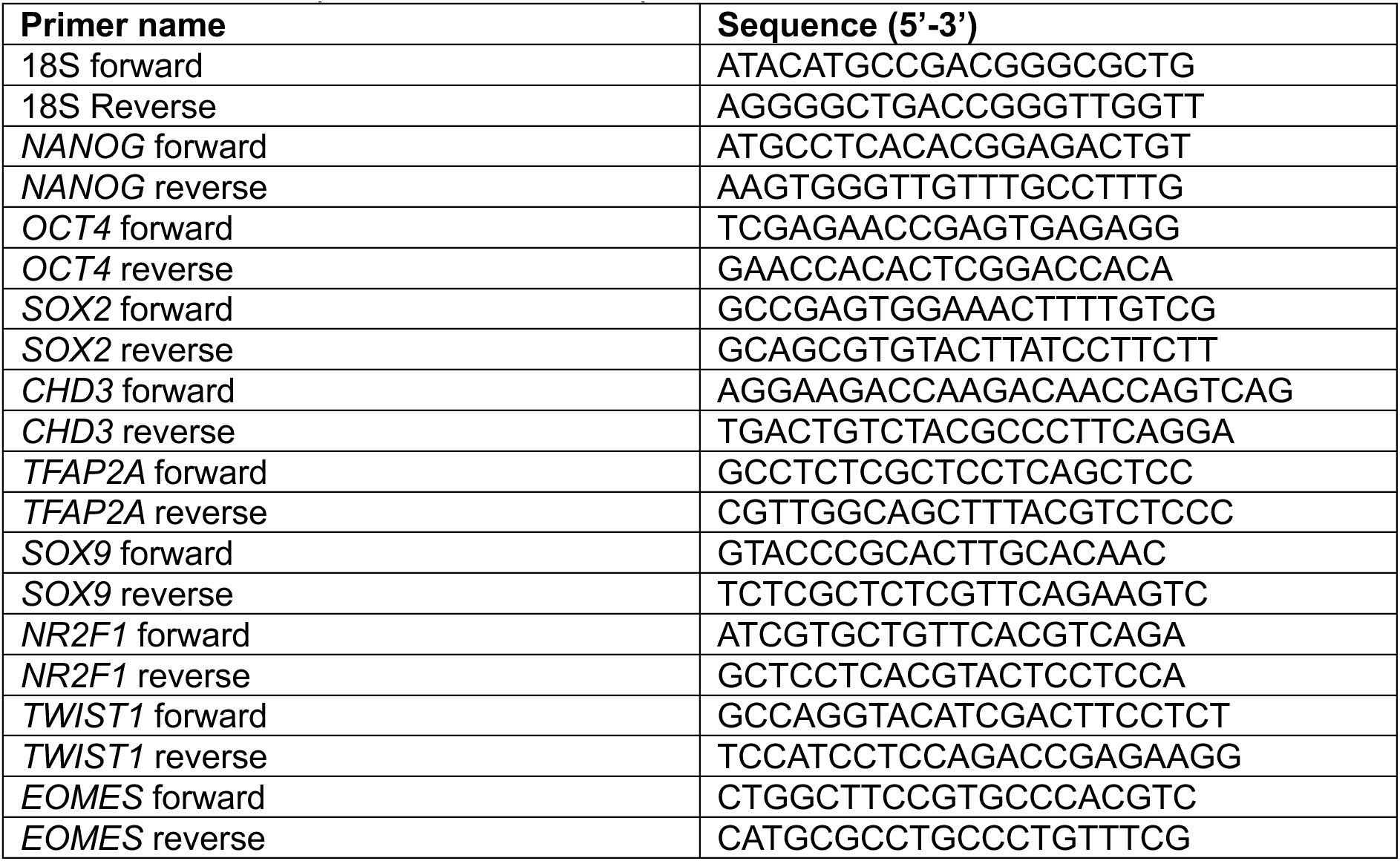
A list of the primers used for RT-qPCR.

### Flow cytometry

Cells were treated with StemPro accutase (Gibco, A1110501) for 5 minutes in order to produce a single-cell suspension. Cells were washed in cold PBS containing 2% fetal bovine serum (FBS) (Gibco, A4736201) and counted using a countess automated cell counter. For each sample and time point, 1×10^6^ cells were resuspended in 100ul PBS-2% FBS and stained with 4ul PE anti-human TRA-1-60-R Antibody (BioLegend, 330609), 2ul APC anti-human SSEA-4 Antibody (BioLegend, 330417) and 500 ng/ml DAPI (4’,6-Diamidino-2-Phenylindole, Dihydrochloride) (BioLegend, 422801). Cells were then incubated protected from light on ice for 15 minutes. Subsequently, cells were filtered into FACS tubes containing 300ul of PBS-2% FBS and flow cytometry was performed using the Agilent NovoCyte Penteon Flow Cytometer at the Sir Alexander Fleming building flow cytometry facility at Imperial College London. The resulting data were analysed using FlowJo software version 10.9.0.

### Immunofluorescence

Cells were plated onto geltrex coated coverslips and fixed using 4% formaldehyde (Fisher Scientific, 10532955) for 15 minutes at 37°C. Samples were then permeabilised with 0.1% Triton-X100 (Merck, 648463) in PBS for 10 minutes at room temperature, before blocking for 1 hour at room temperature in 10% donkey serum (Abcam, AB7475). Samples were incubated overnight at 4°C on a rocker with primary antibodies of interest (table 3) diluted in PBS. Negative controls with only PBS were also set up for each different secondary antibody. Samples were then washed in 0.1% tween 20 (Promega, H5152) in PBS before being stained with the relevant secondary antibodies (table 3) diluted in PBS for 1 hour at 37°C. Samples were washed again in 0.1% tween 20 and stained with 1 ug/ml DAPI (BioLegend, 422801) for 15 minutes at room temperature. Coverslips were mounted using fluorescence mounting medium (Agilent, S302380-2). Stained cells were visualised using the Zeiss Axio Observer inverted microscope in the Facility for Imaging by Light Microscopy (FILM) at Imperial College London.

**Table 3.**
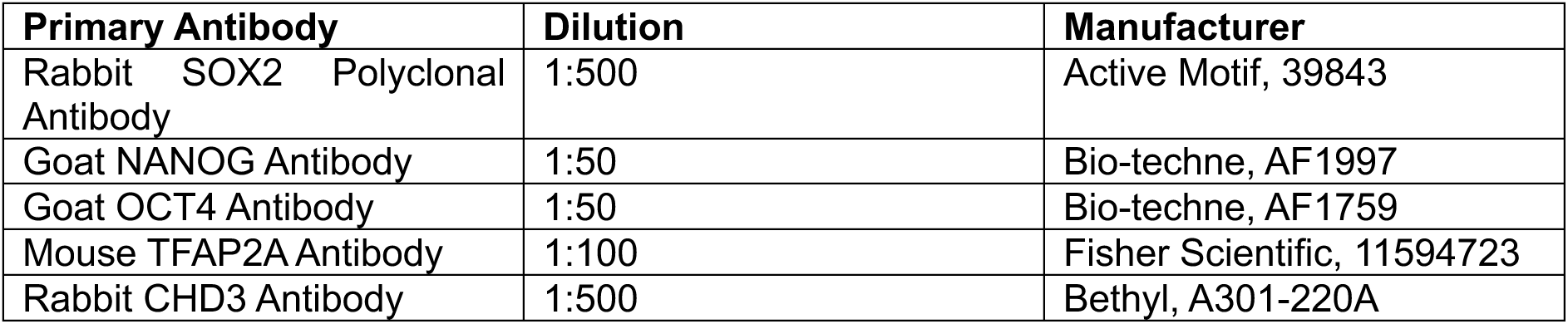
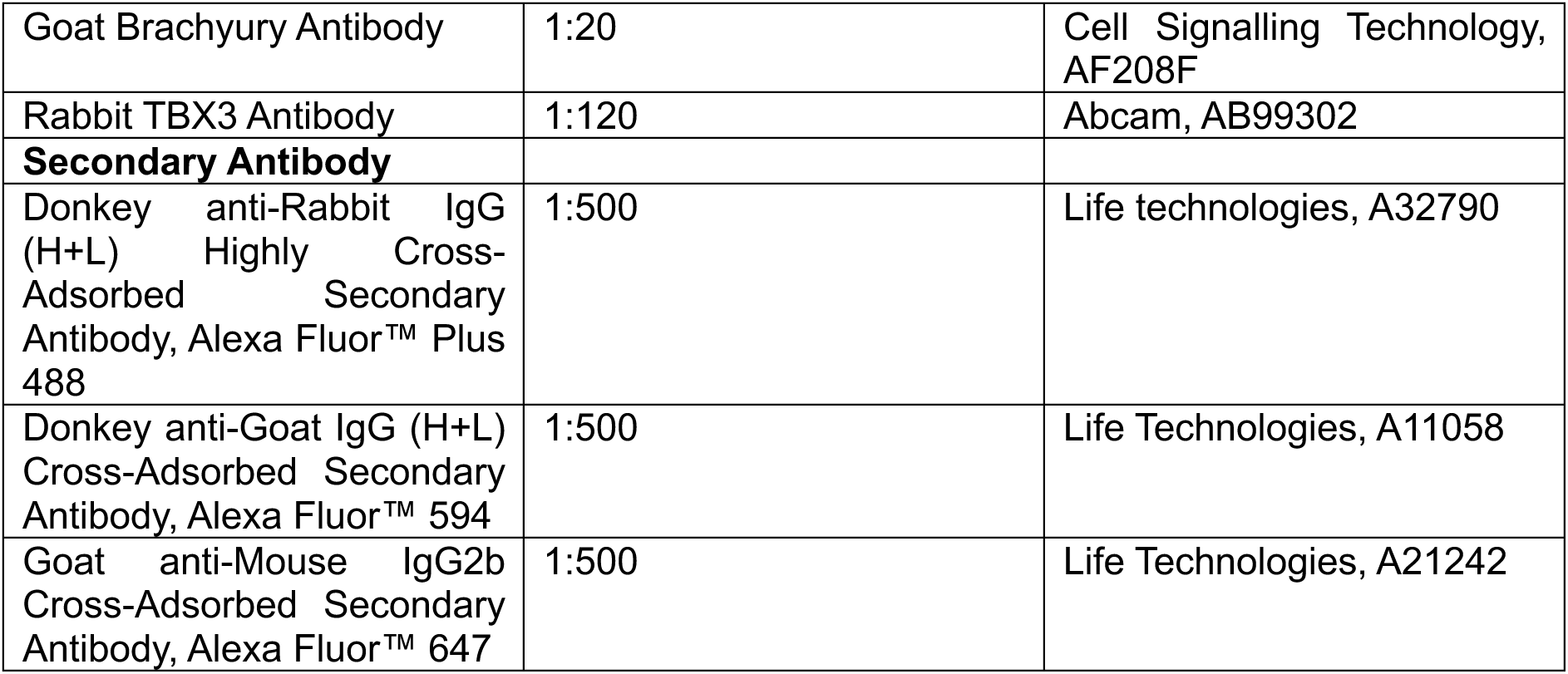
Antibodies used for immunofluorescence.

### Western blot

For the western blot, cells were harvested and washed 3 times in 1X phosphate buffered saline (PBS) and pelleted. Protein was extracted using radioimmunoprecipitation assay (RIPA) buffer (Thermo Scientific, 10230544) with protease inhibitor (PI) and was quantified using the Pierce BCA Protein Assay Kit (Thermo Scientific, 23225). 30ug of each protein sample were loaded onto a Novex Tris-Glycine Mini Protein Gel (Invitrogen, XP04122BOX). Proteins were separated using gel electrophoresis in 1X SDS running buffer and were then transferred to nitrocellulose membrane. The membrane was blocked in Intercept (PBS) Blocking Buffer (LI-COR, 927-70001) for 1 hour. Primary antibodies (table 4) were diluted 1:1000 in Intercept (PBS) Blocking Buffer containing 0.2% Tween 20. The membrane was incubated in the primary antibody dilution on rollers overnight at 4°C. The membrane was then washed 4 times for 5 minutes in 1X PBS containing 0.1% Tween 20 (PBST) on a rocker. The secondary antibody (table 4) was diluted 1:15000 in Intercept (PBS) Blocking Buffer containing 0.2% Tween 20. The membrane was incubated in the secondary antibody protected from light at room temperature for 1 hour on a rocker. The membrane was then washed again in PBST 4 times for 5 minutes on a rocker before being imaged using the LI-COR Odyssey XF imaging system.

**Table 4.** A list of the antibodies used for western blots.

| **Primary Antibody** | **Manufacturer** |
| --- | --- |
| Rabbit anti-CHD3 Antibody | Bethyl Laboratories, A301-220A |
| GAPDH (D16H11) XP Rabbit mAb | Cell Signalling Technology, 5174 |
| **Secondary Antibody** |  |
| IRDye® 800CW Goat anti-Rabbit IgG Secondary Antibody | LI-COR, 926-32211 |

### RNA-sequencing

Cells were pelleted and RNA extraction was performed using the Monarch Total RNA Miniprep Kit (NEB, T2010). RNA was quantified with a nanodrop and the quality was assessed using TapeStation 2200 (Agilent Technologies). Only RNA with a RIN score above 8 was used. RNA libraries were prepared from 1ug of RNA input using the NEBNext Poly(A) mRNA Magnetic Isolation Module (NEB, E7490) and NEBNext Ultra II Directional RNA Library Prep Kit for Illumina (NEB, E7760). Sequencing was carried out by the Wistar Institute to generate 60 bp paired-end reads. Two replicates of clone 1 and one replicate of clone 2 were used at each specified timepoint for all of the three conditions (*CHD3*-WT, *CHD3*-HET-KO and *CHD3*-KO). This corresponded to two biological replicates and a total of three technical replicates per condition.

### RNA-sequencing analysis

First, adapters were removed using TrimGalore!, then reads were mapped and quantified using Kallisto.^77^ Differential gene expression was analysed using DESeq2.^78^ Gene set enrichment analysis was performed using WebGestalt 2019.^79^ Additional statistical analysis was carried out using R (version 4.2.2) and GraphPad Prism (version 10.1.1).

### ATAC-sequencing

ATAC-seq was performed using 50,000 cells per sample. Libraries were prepared using the ATAC-seq Kit (Active Motif, 53150) following the manufacturer’s instructions. Sequencing was carried out by the Wistar Institute to generate 60 bp paired-end reads. One replicate of clone 1 and one replicate of clone 2 were used at each specified timepoint for both conditions (*CHD3*-WT and *CHD3*-KO). This corresponded to two biological replicates per condition.

### ATAC-sequencing analysis

Adapters were removed using TrimGalore! and the reads were then aligned to the hg19 human reference genome using the Burrows-Wheeler Alignment (BWA) tool with the MEM algorithm.^80^ SAMTools^81^ was then used to filter for high quality (MAPQ > 10) reads and to remove PCR duplicates. Peaks were then called using MACS2^82^ with 5% FDR. Consensus peaks (peaks found in all replicates) were identified for each cell line using BEDTools^83^ and these were used for subsequent analyses. ATAC-seq peaks were visualised using UCSC Genome Browser.^84^ Motif anaylsis was performed using HOMER^85^ All further downstream analysis was performed using BEDTools^83^ and deepTools.^86^

### ChIP-sequencing

Two replicates were performed for each condition. For each replicate, 11 million cells were cross-linked using 1% formaldehyde for 5 minutes at room temperature. Cells were then quenched with 125 mM glycine for 5 minutes at room temperature before being washed twice with 1X PBS. The fixed cells were then resuspended in ChIP buffer (150 mM NaCl, 1% Triton X-100, 5 mM EDTA, 10 mM Tris-HCl, 0.5 mM DTT, 0.3% SDS, protease inhibitor) and incubated on ice for 10 minutes. Chromatin was sheared to an average length of 100-1000bp using a Covaris M220 Focused-Ultrasonicator at 5% duty factor for 7 minutes. The chromatin lysate was diluted in SDS-free ChIP buffer. 10ug of antibody was used for both CHD3 (Bethyl Laboratories, A301-220A) and DLX5 (Abcam, AB109737). The antibody was added to at least 5ug of sonicated chromatin along with Dynabeads Protein A (Invitrogen, 10002D) and incubated overnight at 4°C with rotation. The beads were then washed twice with each of the following buffers: Mixed Micelle Buffer (150 mM NaCl, 1% Triton X-100, 0.2% SDS, 20 mM Tris-HCl, 5 mM EDTA, 65% sucrose), Buffer 200 (200 mM NaCl, 1% Triton X-100, 0.1% sodium deoxycholate, 25 mM HEPES, 1 mM EDTA), LiCl detergent wash (250 mM LiCl, 0.5% sodium deoxycholate, 0.5% NP-40, 10 mM Tris-HCl, 1 mM EDTA) and a final wash was performed with cold 0.1X TE. Finally, beads were resuspended in 1X TE containing 1% SDS and incubated at 65°C for 10 min to elute immunocomplexes. The elution was repeated twice, and the samples were incubated overnight at 65°C to reverse cross-linking, along with the input (5% of the starting material). The DNA was digested with 0.5 mg/ml Proteinase K for 1 hour at 65°C and then purified using the ChIP DNA Clean & Concentrator kit (Zymo, D5205) and quantified with the QuantiFluor ONE dsDNA system (Promega, E4871). Barcoded libraries were made with NEBNext Ultra II DNA Library Prep Kit for Illumina (NEB, E7645L) using NEBNext Multiplex Oligos Dual Index Primers for Illumina (NEB, E7600S). Sequencing was carried out by the Wistar Institute to generate 60 bp paired-end reads or by Novogene to generate 150 bp paired-end reads. Clones 1 and 2 were used as biological replicates.

### ChIP-sequencing analysis

Adapters were removed with TrimGalore! and the sequences were aligned to the reference genome hg19, using Burrows-Wheeler Alignment tool, with the MEM algorithm.^80^ Uniquely mapping aligned reads were filtered based on mapping quality (MAPQ > 10) to restrict our analysis to higher quality and uniquely mapped reads, and PCR duplicates were removed. HOMER^85^ was used to call peaks using the default parameters at 5% FDR. All statistical analyses were performed using BEDTools^83^, deepTools^86^, R (version 4.2.2) and GraphPad Prism (version 10.1.1).

### Trilineage differentiation

iPSCs were differentiated into the three germ layers (endoderm, ectoderm and mesoderm) using the Human Pluripotent Stem Cell Functional Identification Kit (R&D Systems, SC027B). Expression of relevant markers was then assessed using immunofluorescence with 10 ug/ml of the antibodies provided in the kit (Goat anti-human SOX17, Goat anti-human Otx2 and Goat anti-human Brachyury).

**Table 3.** Antibodies used for trilineage immunofluorescence.

| Primary Antibody | Dilution/concentration | Manufacturer |
| --- | --- | --- |
| Goat anti-human SOX17 | 10 ug/ml | R&D systems, 963121 |
| Goat anti-human Otx2 | 10 ug/ml | R&D systems, 963273 |
| Goat anti-human Brachyury | 10 ug/ml | R&D systems, 963427 |
| Secondary Antibody |  |  |
| Donkey anti-Goat IgG (H+L) Cross-Adsorbed Secondary Antibody, Alexa Fluor™ 594 | 1:500 | Life Technologies, A11058 |

## Notes

### Competing Interest Statement

The authors have declared no competing interest.

