## Supplementary Figures for "CHD3 regulates BMP signalling response during cranial neural crest cell specification"

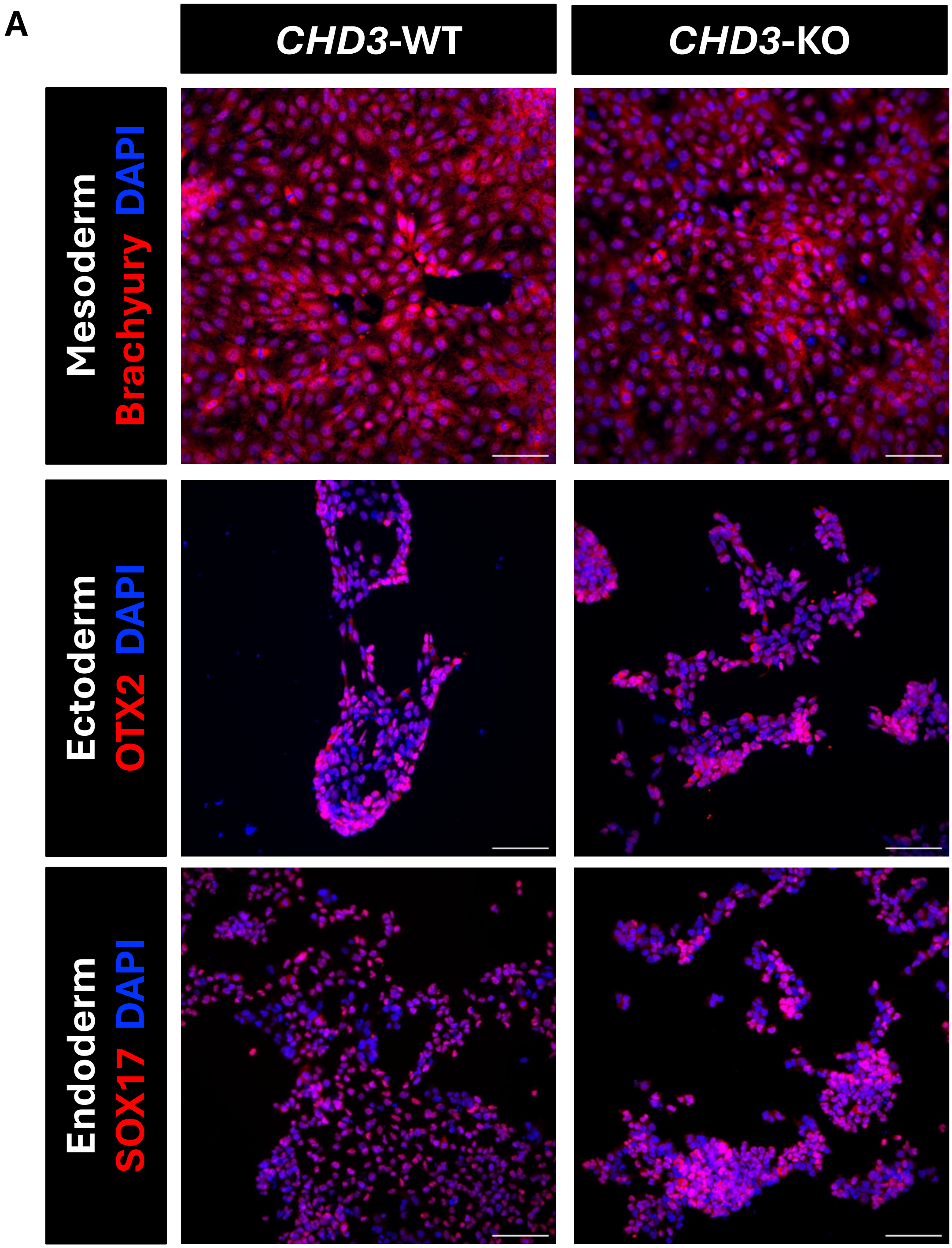


**Figure S1. Validation of the iPSC lines with a trilineage assay.** (A) Immunofluorescence for mesoderm marker brachyury, ectoderm marker OTX2 and endoderm marker SOX17 in *CHD3*-WT and *CHD3*-KO following differentiation into the three respective germ layers. Scale bar: 50 µm.

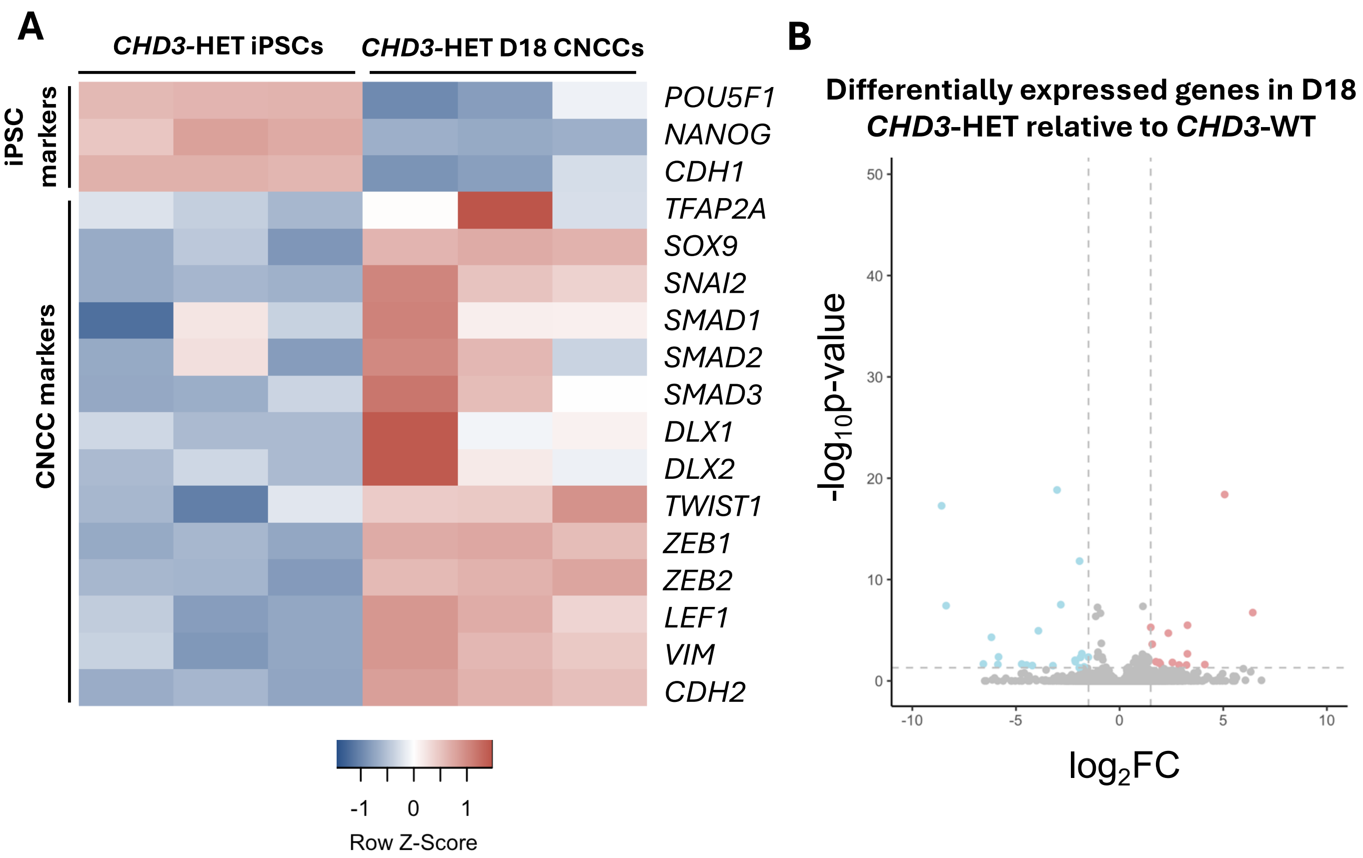


**Figure S2. Characterisation of *CHD3*-HET D18 CNCCs.** (A) Heatmap displaying expression of key pluripotency markers and CNCC markers in *CHD3*-HET iPSCs and *CHD3*-HET D18 CNCCs. (B) Volcano plot of differentially expressed genes in *CHD3*-HET comparative to *CHD3*-WT in D18 CNCCs. Blue dots represent downregulated genes with p-adj < 0.05 and log2FoldChange < -1.5. Red dots represent upregulated genes with p-adj < 0.05 and log2FoldChange > 1.5.

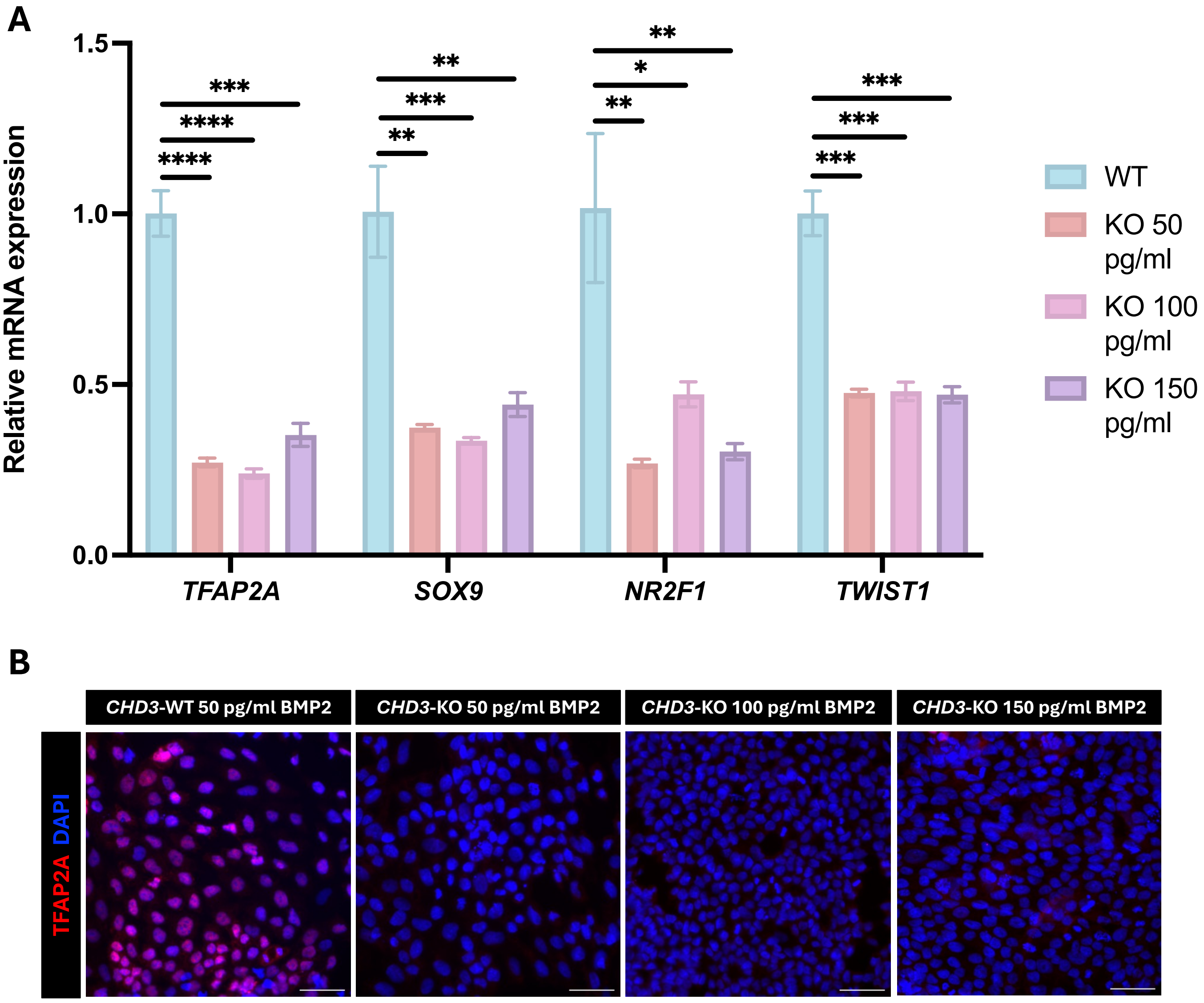


**Figure S3. Increasing BMP2 dosage does not rescue CNCC marker expression in *CHD3*-KO D18 CNCCs.** (A) RT-qPCR assessing the relative expression levels of CNCC markers (*TFAP2A*, *SOX9*, *NR2F1* and *TWIST*) between *CHD3*-WT (WT) and *CHD3*-KO (KO) D18 CNCCs provided with either the standard 50 pg/ml or an increased concentration (100 pg/ml or 150 pg/ml) of BMP2. *CHD3*-WT was given the standard 3 μM CHIRON and *CHD3*-KO was given 1 μM CHIRON. Differences between conditions were assessed using unpaired student's t-test. *=p<0.05, **=p<0.01, ***=p<0.001, ****=p<0.0001. (B) Immunofluorescence for the CNCC marker TFAP2A in CHD3-WT and CHD3-KO D18 CNCCs provided with either the standard 50 pg/ml or an increased concentration (100 pg/ml or 150 pg/ml) of BMP2. *CHD3*-WT was given the standard 3 μM CHIRON and *CHD3*-KO was given 1 μM CHIRON. Scale bar: 50 μm.
